# p120 RasGAP and ZO-2 are essential for Hippo signaling and tumor suppressor function mediated by p190A RhoGAP

**DOI:** 10.1101/2023.05.22.541483

**Authors:** Hanyue Ouyang, Wangji Li, Steen H. Hansen

## Abstract

*ARHGAP35*, which encodes p190A RhoGAP (p190A), is a major cancer gene. p190A is a tumor suppressor that activates the Hippo pathway. p190A was originally cloned via direct binding to p120 RasGAP (RasGAP). Here, we determine that a novel interaction of p190A with the tight junction-associated protein ZO-2 is dependent on RasGAP. We establish that both RasGAP and ZO-2 are necessary for p190A to activate LATS kinases, elicit mesenchymal-to-epithelial transition, promote contact inhibition of cell proliferation and suppress tumorigenesis. Moreover, RasGAP and ZO-2 are required for transcriptional modulation by p190A. Finally, we demonstrate that low *ARHGAP35* expression is associated with shorter survival in patients with high, but not low, transcript levels of *TJP2* encoding ZO-2. Hence, we define a tumor suppressor interactome of p190A that includes ZO-2, an established constituent of the Hippo pathway, and RasGAP, which despite strong association with Ras signaling, is essential for p190A to activate LATS kinases.

## Background

Major GWAS studies published almost a decade ago identified *ARHGAP35* as one of the 30-40 most significantly mutated genes in tumor samples ^1,2^. At that time, it was further determined that the region on chromosome 19, where the *ARHGAP35* gene is located, ranks among the most frequently lost in cancer ^3^. Since then, there has been a steady trickle of additional GWAS studies associating *ARHGAP35* alterations with human cancer ^4–14^. Yet, this body of work has been unable to assign a role for *ARHGAP35* in cancer beyond noting that the spectrum of alterations is suggestive of a tumor suppressor function. *ARHGAP35* encodes p190A RhoGAP (p190A), an enzyme that promotes GTP hydrolysis on Rho and Rac GTPases ^15,16^. However, p190A is a large and complex protein with a GTPase domain, four FF motifs, two pseudo-GAP domains and an active GAP domain, in addition to sequences of unknown function. These domains modulate GAP activity but also exert scaffolding functions, as exemplified by the direct binding of Rnd proteins and sequestration of the TFII-I transcription factor ^17,18^.

Rooted in GWAS data, we have relied on unbiased approaches to elucidate a role for p190A in epithelial oncogenesis, because *ARHGAP35* alterations are predominantly found in carcinomas (gdc.cancer.gov). Our initial efforts have led to the discovery that p190A promotes contact inhibition of cell proliferation (CIP) via modulation of both the canonical Hippo pathway and mechano-transduction to repress the activity of the proto-oncogenic transcriptional co-activator YAP ^19^. Moreover, we determined that restoring p190A expression in cancer cells with defined *ARHGAP35* alteration induces expression of *CDH1* encoding E-cadherin to establish a feed-forward loop that activates LATS kinases and promotes mesenchymal-to-epithelial transition (MET). We furthermore provided the first direct demonstration of p190A functioning as a tumor suppressor in vivo and determined that recurrent *ARHGAP35* mutations in human tumor samples exhibit loss of function ^20^.

A role for p190 proteins in promoting GTP-hydrolysis of Rho proteins is well established ^21^. In contrast, it is unclear how p190A might activate the Hippo pathway. In this work, we have used mass spectrometry to identify p190A-interacting proteins that may account for the capacity of p190A to activate LATS kinases. The most frequently interacting proteins identified were RasGAP and ZO-1/2 encoded by *RASA1* and *TJP1/2*, respectively. Here, we determine that p190A binds ZO-1/2 in a RasGAP-dependent manner, and that interactions with RasGAP and ZO-2 are obligatory for p190A to activate LATS kinases, elicit MET, induce CIP and suppress tumorigenesis. Moreover, we establish that RasGAP and ZO-2 play essential role in transcriptional alterations elicited by expression of p190A. Finally, we demonstrate that expression of *ARHGAP35* in human tumors correlates with Hippo signaling, and that the impact of *ARHGAP35* alteration on survival of patients with lung adenocarcinoma (LUAD) is influenced by *TJP1* and *TJP2* transcript levels. Collectively, these findings define a tumor suppressor interactome of p190A consisting of RasGAP ^22^, a negative regulator of Ras signaling, and ZO-2, a tight junction constituent and modulator of YAP/TAZ transcriptional co-activators ^23^.

## Results

### Direct binding to RasGAP is required for p190A to activate LATS kinases, elicit MET and promote CIP

Our published studies have established a pivotal role for p190A in activation of LATS kinases ^19,20^. However, the mechanism by which p190A exerts this function is not well understood. In this work, we tested the hypothesis that p190A might bind to upstream constituents of the canonical Hippo pathway. To this end, we exploited the fact that expression of Myc-tagged p190A in NSCLC NCI-H661 cells, which harbor defined *ARHGAP35* loss-of-function alterations, activates LATS kinases to suppress oncogenic capacities ^20^. Hence, in these H661-p190A cells, we have established that the link between p190A and the canonical Hippo pathway is intact. We therefore immunoprecipitated p190A with anti-Myc 9E10 antibody from control and H661-p190A cells, and subjected the samples for mass spectrometry analysis. Two classes of co-immunoprecipitating proteins were identified more frequently than others (**Figure S1**). The first was RasGAP, which was reassuring, as p190A initially was identified cloned via its direct binding to RasGAP^15,24^. The second was the tight junction-associated Zonula Occludens proteins ZO-1 and ZO-2 ^25,26^.

We initially focused on the interaction with RasGAP, which is well-established to augment the RhoGAP activity of p190A and to promote directional motility ^27,28^. However, in cancer biology, RasGAP is primarily implicated in Ras signaling and is invariably assigned this role in GWAS ^22^. We considered the intriguing possibility that RasGAP might impact the canonical Hippo pathway via p190A. To this end, we knocked out *RASA1* encoding RasGAP from H661 cells using CRISPR/Cas9 technology. Strikingly, p190A was unable to activate LATS kinases in cells depleted of RasGAP (**Figure 1A and Figure 1B**). Equally, the capacity of p190A to induce CIP, which is mediated by the Hippo pathway, was significantly perturbed in H661-p190A *RASA1* cells (**Figure 1A and Figure 1B**). Next, we tested whether direct interaction between RasGAP and p190A is necessary for these effects. The p190A-RasGAP interaction is mediated via FAK/Src phosphorylation of Tyr1087 and Tyr1105 in p190A, which creates docking sites for two SH2 domains in RasGAP that are separated by an SH3 domain (**Figure S2A**) ^29–31^. This interaction is abrogated by mutation of Tyr1087 and Tyr1105 in p190A to phenylalanines. Here, we abbreviate this double-mutant p190A(Y2F). In contrast to wild type p190A, expression p190A(Y2F) in H661 cells failed to activate LATS kinases or induce CIP, as determined by western blotting of total cell lysates to detect pLATS1(S909) and cyclin A, respectively (**Figure 1C and Figure 1D**). Of note, the anti-pLATS1(S909) antibody also reacts with pLATS2(S872), thus precluding distinction between the two paralogs. The requirement for p190A-RasGAP complex formation to induce CIP was further evidenced by quantification of cell number after seeding cells at sparse density. H661 cells expressing p190A(wt) reached a saturation density of ∼3 million cells in a 35-mm dish or ∼3×10^5^ cells/cm^2^. In contrast, control cells or cells expressing p190A(Y2F), as well a H661-p190A cells with *RASA1*-ko grew to significantly higher densities without the growth curve leveling out (**Figure 1E**). This difference in growth characteristics was also evidenced by confocal and phase microscopy. Only cells reconstituted with p190A(wt) grew in cobblestone-like monolayers, while control cells or cells with perturbed p190A-RasGAP complex formation exhibited cell multilayering (**Figure 1F**).

**Figure 1:**
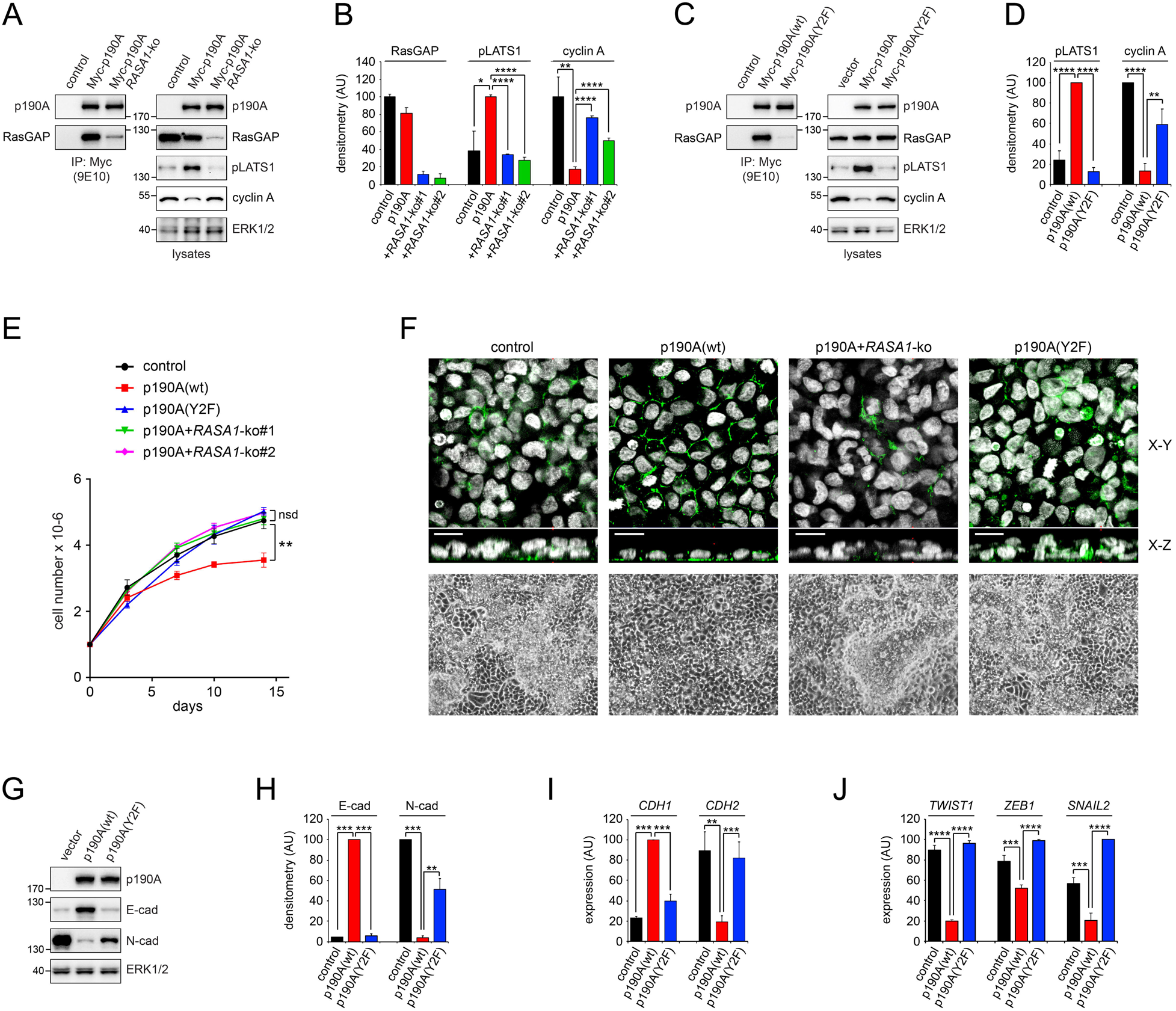
Interaction of p190A with RasGAP is necessary for p190A to activate of LATS1 kinase, promote CIP and elicit MET. (**A**) Western blots of co-IPs and corresponding whole cell lysates to detect p190A, RasGAP, pLATS1, cyclin A and ERK1/2 in control H661 cells, as well as H661-p190A cells w/wo knockout of *RASA1*. (**B**) Quantification by densitometry of RasGAP, pLATS1, and cyclin A levels detected by western blotting as shown in (**A**). Data are presented as mean ± SD (n=3). (**C**) Western blots of co-IPs and corresponding whole cell lysates to detect p190A, RasGAP, pLATS1, cyclin A and ERK1/2 in control cells, as well as H661 cells reconstituted with either p190A(wt) or p190A(Y2F) defective in RasGAP-binding. (**D**) Quantification by densitometry of RasGAP, pLATS1, and cyclin A levels detected by western blotting as shown in (**C**). Data are presented as mean ± SD (n=3). (**E**) Growth curves for control and H661 cells reconstituted with p190A(wt) or p190A(Y2F), with or without knockout of *RASA1*. 1×10^6^ cells were seeded per well of a 6-well plate and propagated for the number of days indicated with change of medium every 2 days. Cell number was quantified manually. Data are presented as mean ± SD (n=3). Statistical testing was performed by pairwise Student’s *t* test as indicated by brackets. **p<0.025; nsd: not significantly different. (**F**) The p190A-RasGAP interaction is necessary for the capacity of p190A to restore growth of H661 cells in monolayers. Top panels show confocal microscopy of control cells, as well as H661 cells with expression of p190A(wt) or p190A(Y2F) and with or without *RASA1* knockout. Scale bars correspond to 10 µm. The bottom panels show phase images for the same conditions. (**G**) Western blots of whole cell lysates to detect p190A, E-cadherin, N-cadherin and ERK1/2 in control cells, as well as H661 reconstituted with either wild type p190A or p190A(Y2F). (**H**) Quantification by densitometry of E-cadherin, and N-cadherin levels detected by western blotting as shown in (**g**). Data are presented as mean ± SD (n=3). (**I**) Transcript levels for *CDH1* and *CDH2*, as determined by qPCR. Data are presented as mean ± SD (n=4). (**J**) Transcript levels for *TWIST1*, *ZEB1* and *SNAIL2*, as determined by qPCR. Data are presented as mean ± SD (n=4). All statistical testing for data presented in this figure was performed using pairwise Student’s *t* test as indicated by brackets. *p<0.05; **p<0.025; ***p<0.01; ****p<0.001.

We furthermore tested the requirement for the p190A-RasGAP interaction to elicit MET. Unlike p190A(wt), expression of p190A(Y2F) did not promote an N- to E-cadherin switch (**Figure 1G-I**). Moreover, p190A(Y2F) failed to modulate expression of the *TWIST1*, *ZEB1* and *SNAIL2* genes that we previously have shown are suppressed by expression of p190A(wt) (**Figure 1J**) ^20^. Thus, formation of a p190A-RasGAP complex is required for LATS activation, to elicit MET and promote CIP in H661 cells. Of note, we obtained very similar results with NSCLC NCI-H226 cells, thereby suggesting general relevance of our findings (**Figure S2B-F**).

### Interaction with RasGAP is necessary for the tumor suppressor function of p190A

Next, we tested a role for the p190A-RasGAP interaction in tumor suppressor function using a xenograft model that we previously established for p190A ^20^. We injected 5×10^6^ control H661 cells or cells expressing either p190A(wt) or p190A(Y2F) subcutaneously into the flank of nude mice. As previously reported ^20^, mice injected with H661-p190A(wt) cells exhibited small tumors that rapidly regressed (**Figure 2A**). In contrast, the cohorts of mice injected with control cells or p190A(Y2F) cells showed aggressive tumor growth, which necessitated euthanasia after 4-10 weeks for all control mice and two thirds of p190A(Y2F) mice (**Figure 2A and Figure 2B**). The p190A(Y2F) tumors were slightly smaller than the control tumors but the difference was not statistically significant. However, the difference was reflected in survival (**Figure 2B**). There were no apparent histological features to distinguish tumors formed by control and p190A(Y2F) cells (**Figure S3A**), with the exception that we detected staining for p190A in p190A(Y2F) tumors but not in control tumors (**Figure S3B**). Likewise, we did not observe overt differences in Ki-67 staining between control and p190A(Y2F) tumors (**Figure S3C**), which was verified by quantification (**Figure S3D**).

**Figure 2:**
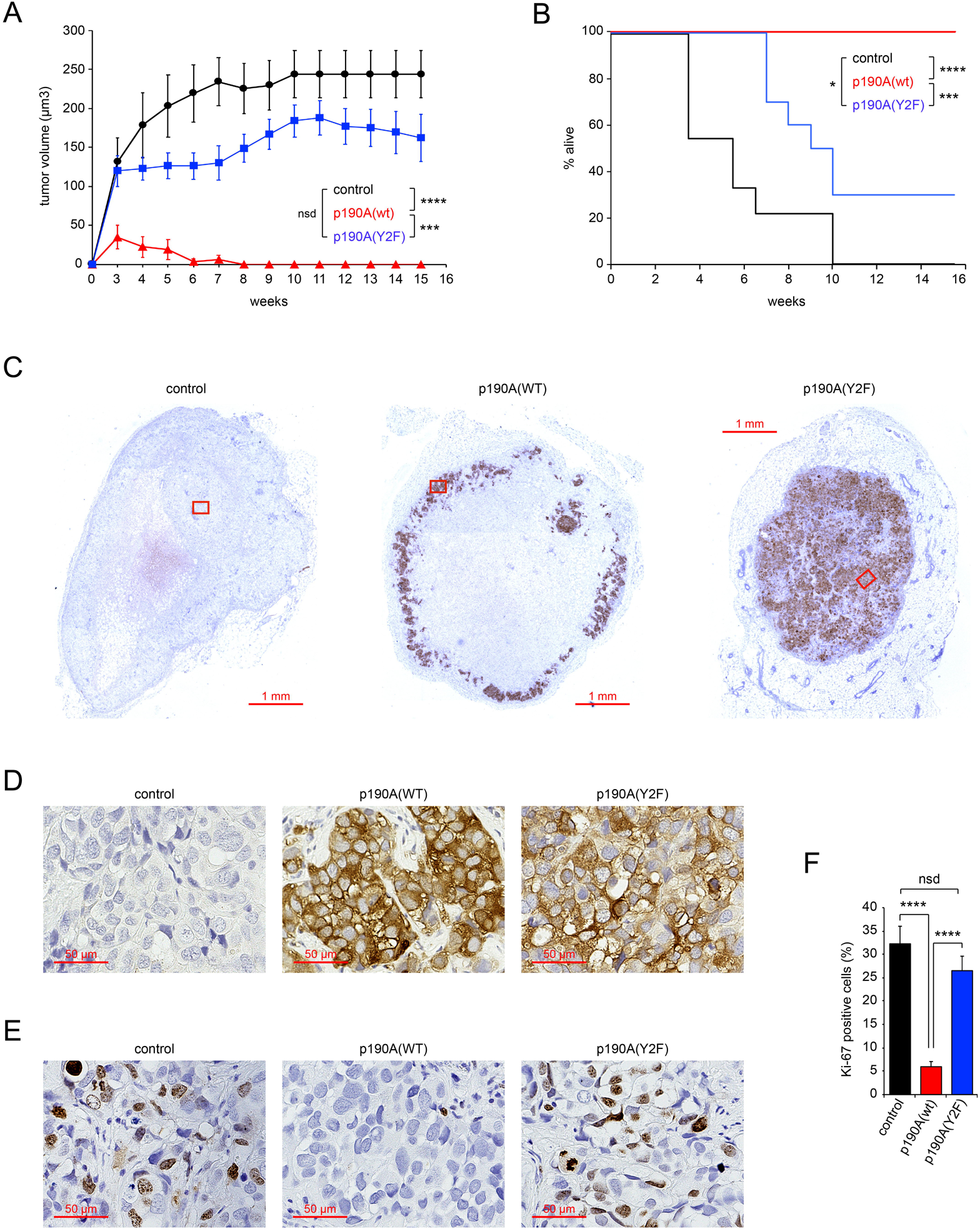
Interaction of p190A with RasGAP is required for the tumor suppressor function of p190A. (**A**) Cumulative xenograft tumor volume in mice injected with control cells (n=9) or H661 cells expressing p190A(wt) (n=10) or p190A(Y2F) (n=10). Statistical testing was performed using pairwise Student’s *t* test as indicated by brackets. ***p<0.01; ****p<0.001; nsd: not significantly different. (**B**) Kaplan-Meier survival plot for mice the cohort of mice described in (**A**). Statistical testing was performed using pairwise log-rank test as indicated by brackets. *p<0.05; ***p<0.01; ****p<0.001. (**C**) Immunohistochemistry to detect p190A in tumors 3 weeks after injection of control cells or H661 cells expressing p190A(wt) or p190A(Y2F). (**D**) 20x magnifications of the areas contained within the red boxes in (**C**). (**E**) Immunohistochemistry to detect Ki-67 positive cells in tumors 3 weeks after injection of control cells or H661 cells expressing p190A(wt) or p190A(Y2F). (**F**) Quantification of Ki-67 positive cells from (**E**). Data are presented as mean ± SEM with n=5 mice and >600 nuclei scored per condition. Statistical testing was performed using pairwise Student’s *t* test as indicated by brackets. ****p<0.001; nsd: not significantly different.

In order to compare tumors formed from cells expressing p190A(Y2F) to p190A(wt), we initiated a second cohort where all mice were sacked three weeks after injection of control, p190A(wt) or p190A(Y2F) cells. This analysis revealed that the center of p190A(wt) tumors was mostly comprised by debris with the tumor cells lining the periphery (**Figure 2C**). In contrast, control and p190A(Y2F) tumors were typically populated by cells both peripherally and internally (**Figure 2C**). We did not observe major differences in cell morphology between control, p190A(wt) or p190A(Y2F) tumors in this cohort (**Figure S3E**). Moreover, p190A protein was readily detected by IHC in both p190A(wt) and p190A(Y2F) tumors, but absent from tumors formed from control cells (**Figure 2D**). Finally, staining for Ki-67 revealed that while p190A(wt) cells were quiescent, p190A(Y2F) cells were actively proliferating, to equal extent as control cells (**Figure 2E and Figure 2F**). Taken together, these results demonstrate that direct interaction with RasGAP is necessary for the tumor suppressor function of p190A.

### ZO-1 and ZO-2 bind p190A in a RasGAP-dependent manner and dictate p190A localization

We then turned our attention to the novel interaction between p190A and ZO-1/2 identified by mass spectrometry. First, we validated the interaction. Using anti-Myc 9E10 antibody coupled to agarose beads, we performed IP from control and H661-p190A cells. Following SDS-PAGE of precipitated proteins and western blotting, we readily detected ZO-1 and ZO-2 in IPs from H661-p190A cells, but not control cells (**Figure 3A**). Second, we verified that the interaction occurs between endogenously expressed p190A and ZO-1/2 proteins in MDA-MB-231 cells (**Figure 3B**). Third, we tested if binding of p190A to RasGAP was necessary for the interaction with ZO-1/2 in H661 cells. Indeed, perturbation of the interaction between p190A and RasGAP, either by knockout of *RASA1* from p190A(wt) cells or by expression of p190A(Y2F), obliterated the interaction between p190A and ZO-1/2 (**Figure 3C**). In contrast, knockout of *TJP2* from H661-p190A cells did not impact the binding of RasGAP to p190A (**Figure 3D**). We did not test a role for ZO-1 in the p190A-RasGAP interaction.

**Figure 3:**
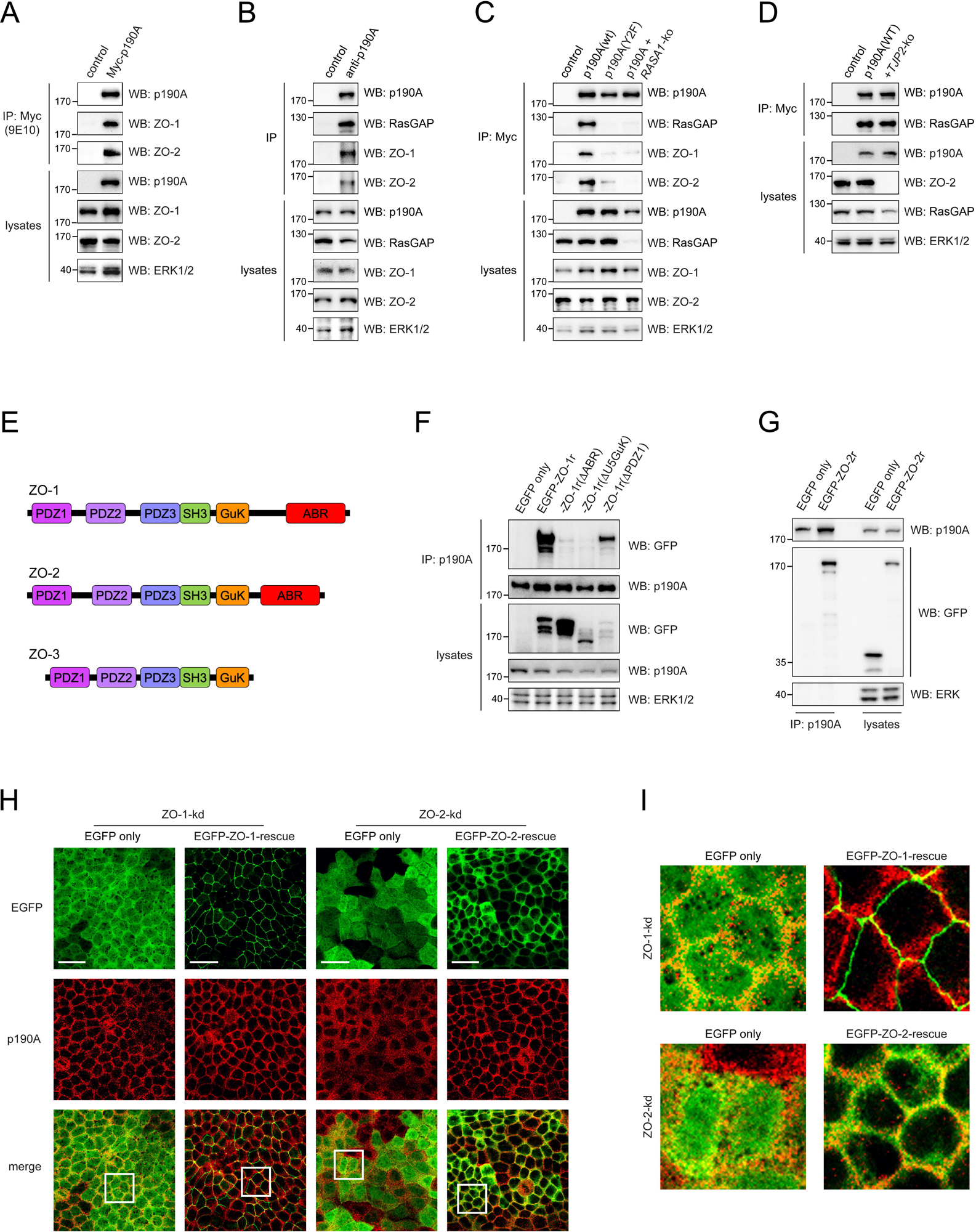
ZO-1/2 bind p190A in a RasGAP-dependent manner and recruits p190A to the plasma membrane. (**A**) Validation of interaction between p190A and ZO-1/2 detected by mass spec. Confluent cultures of control cells and H661 expressing Myc-tagged p190A cells were lysed in GLB and processed for immunoprecipitation using mouse monoclonal 9E10 anti-Myc epitope antibody immobilized on agarose beads followed by western blotting to detect p190A, ZO-1, ZO-2 and ERK1/2. (**B**) Confluent cultures of MDA-MB-231 cells were processed for immunoprecipitation with rabbit polyclonal anti-p190A antibody or without antibody followed by protein A/G sepharose beads. Next, p190A, RasGAP, ZO-1, ZO-2 and ERK1/2 were detected in immunoprecipitates and lysates by western blotting. (**C**) ZO-1 and ZO-2 bind p190A in a RasGAP-dependent manner. Confluent cultures of control cells or H661 cells expressing Myc-tagged p190A(wt) or p190A(Y2F) w/wo RASA1-ko were processed for immunoprecipitation with 9E10 antibody followed by western blotting to detect p190A, RasGAP, ZO-1, ZO-2 and ERK1/2. (**D**) Binding of p190A to RasGAP is independent of ZO-2. Confluent cultures of control cells or H661 cells expressing Myc-tagged p190A w/wo *TJP2*-ko were processed for immunoprecipitation with 9E10 antibody followed by western blotting to detect p190A, RasGAP, ZO-2 and ERK1/2. (**E, F**) The U5GuK and ABR domains of ZO-1 are necessary for the interaction with p190A. (**E**) Cartoon depicting domain structure of Zonula Occludens proteins ZO-1, ZO-2 and ZO-3. (**F**) MDCK II cells depleted of endogenous ZO-1 and expressing exogenous EGFP only or knockdown-resistant EGFP-tagged full-length ZO-1 or ZO-1 deletion mutant missing the ABR, U5GuK, or PDZ1 domains were processed for immunoprecipitation with either polyclonal rabbit anti-p190A antibody followed by western blotting to detect GFP, p190A and ERK1/2. (**G**) Interaction of ZO-2 with p190A in MDCK cells. MDCK II cells depleted of endogenous ZO-2 and expressing exogenous EGFP only or knockdown-resistant EGFP-tagged full-length ZO-2 were processed for immunoprecipitation with polyclonal p190A antibody followed by western blotting to detect GFP and p190A. (**H**) Immunofluorescence to detect p190A (red) in MDCK II cell depleted of ZO-1 or ZO-2 expressing either EGFP alone or knockdown-resistant EGFP-tagged full length ZO-1 or ZO-2 as indicated in the figure. Scale bars correspond to 10 µm. (**I**) 5x magnification of the boxed areas in (**H**).

Next, we probed the requirement for defined motifs in Zonula Occludens proteins for the interaction with p190A (**Figure 3E**). MDCK cells with constitutive knockdown of ZO-1 and conditional expression of knockdown-resistant EGFP-tagged full length ZO-1 or deletion mutants have been engineered and described by others ^32^. From each of these cell lines, we performed IP with anti-p190A antibody followed by SDS-PAGE to detect relevant proteins by western blotting. The results shown in **Figure 3F** demonstrate that the carboxy-terminal GuK and ABR regions of ZO-1 are necessary for the interaction with p190A, while the N-terminal PDZ1 domain is dispensable. We moreover used this approach to determine that p190A binds to EGFP-tagged ZO-2 (**Figure 3G**), thus further supporting the data reported above. We were unable to IP ZO-3 with p190A, which is unsurprising, as ZO-3 lacks an ABR (**Figure 3E**).

We have previously demonstrated that a fraction of p190A in MDCK cells is associated with lateral membranes engaged in cell-cell contact ^19^. Next, we examined if ZO-1/2 mediates recruitment of p190A to these sites by confocal microscopy. In cells depleted of ZO-1, p190A showed membrane localization that was enhanced by expression of EGFP-tagged ZO-1 (**Figure 3H and Figure 3I**). Strikingly, in cells depleted of ZO-2, p190A was localized in puncta scattered diffusely throughout the cytoplasm. Upon, rescue with EGFP-tagged ZO-2, p190A showed intense staining of membranes engaged in cell-cell contact. Thus, ZO-1 and ZO-2 promote targeting of p190A to lateral membranes (**Figure 3H and Figure 3I**). Attempts to determine if ZO-1/2 control the subcellular localization of p190A in H226 and H661 cells lines have thus far been unsuccessful due to low intensity staining. However, the data obtained with MDCK cells establish that ZO-1/2 not only bind to p190A but also can dictate its subcellular localization.

### Interaction with ZO-2 is necessary for p190A to activate LATS kinases, elicit MET, induce CIP and suppress tumorigenesis

To test whether ZO-1/2 are required for tumor suppressor functions of p190A, we generated control and H661-p190A(wt) cells with knockout of *TJP1* or *TJP2*, encoding ZO-1 and ZO-2, respectively (**Figure 4A, Figure 4B and Figure S4A**). To this end, we used a total of two sgRNAs targeting sequences unique to *TJP1* and four sgRNAs targeting sequences unique to *TJP2*. Of note, knockout of *TJP1* did not affect ZO-2 expression (**Figure 4A**). In contrast, knockout of *TJP2* reduced ZO-1 levels (**Figure 4B**). The results of functional analyses revealed remarkable differences between *TJP1* and *TJP2* knockouts. While *TJP1* knockout was largely innocuous (**Figure 4A**), knockout of *TJP2* abolished the capacity of p190A to activate LATS1, induce E-cadherin expression and suppress cyclin A levels (**Figure 4B, Figure 4C and Figure S4A**). Quantification of *CDH1* and *CDH2* transcripts further established that ZO-2 is essential for the N- to E-cadherin switch elicited by expression of p190A (**Figure 4D**). Accordingly, knockout of *TJP2* to a greater extent than knockout of *TJP1* abrogated the suppression of *TWIST1* and *ZEB1* transcripts observed upon restoring p190A expression in H661 cells (**Figure S4B**). Furthermore, in agreement with the effects on cyclin A levels, cell proliferation assays demonstrated that *TJP2*, but not *TJP1*, is obligatory for p190A to induce CIP in H661 cells (**Figure 4E**). Moreover, knockout of *TJP2*, but not *TJP1*, elicited cell multilayering of H661-p190A cells indistinguishable from control cells (**Figure 4F**). Collectively, these results demonstrate that ZO-2 is required for p190A to activate LATS kinases, promote MET and induce CIP. However, we cannot formally exclude the possibility that partial depletion of ZO-1 contributes to the phenotype of *TJP2* knockout cells.

**Figure 4:**
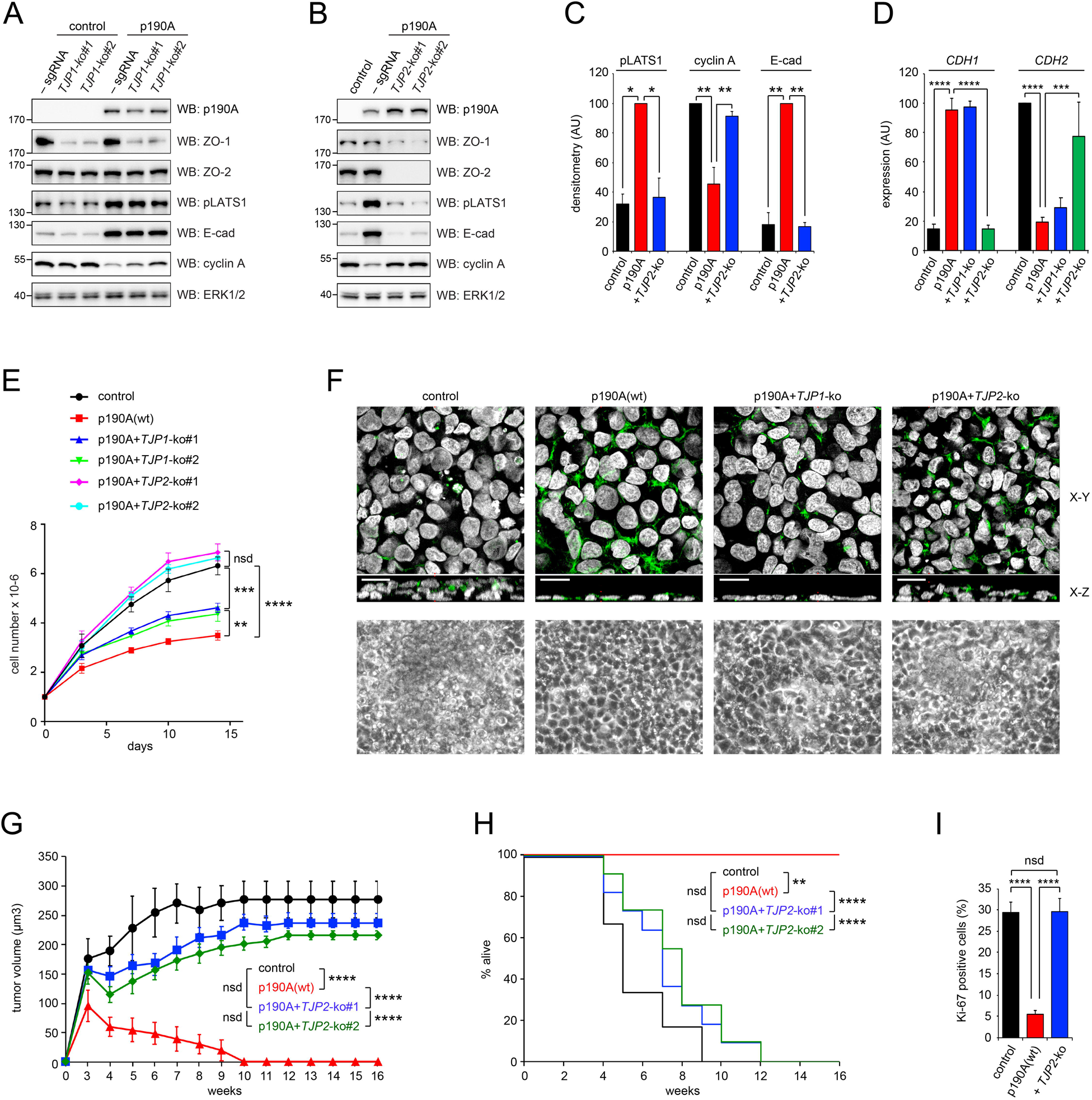
ZO-2 is necessary for p190A to activate LATS1 kinase, promote CIP, elicit MET and suppress tumorigenesis of H661 cells in nude mice. (**A**) Western blots of whole cell lysates to detect p190A, ZO-1, ZO-2, pLATS1, E-cadherin, cyclin A and ERK1/2 in control cells and H661-p190A cells w/wo knockout of *TJP1*. (**B**) Western blots of whole cell lysates to detect p190A, ZO-1, ZO-2, pLATS1, E-cadherin, cyclin A and ERK1/2 in control cells and H661-p190A cells w/wo knockout of *TJP2*. (**C**) Quantification by densitometry of pLATS1, cyclin A and E-cadherin levels detected by western blotting as shown in (**B**). Statistical testing was performed using pairwise Student’s *t* test as indicated by brackets. *p<0.05; **p<0.025. (**D**) Transcript levels for *CDH1* and *CDH2*, as determined by qPCR. Data are presented as mean ± SD (n=6). Statistical testing was performed using pairwise Student’s *t* test as indicated by brackets. ***p<0.01; **p<0.001. (**E**) Growth curves for control and H661-p190A cells with or without knockout of *TJP1 or TJP2*. 1×10^6^ cells were seeded per well of a 6-well plate and propagated for the number of days indicated with change of medium every 2 days. Cell number was quantified manually. Data are presented as mean ± SD (n=3). Statistical testing was performed by pairwise Student’s *t* test as indicated by brackets. **p<0.025; ***p<0.01; ****p<0.001; nsd: not significantly different. (**F**) *TJP2* is necessary for p190A to restore growth of H661 cells in monolayers. Top panels show confocal microscopy of control cells as well as H661 cells with expression of p190A w/wo *TJP1* or *TJP2* knockout. Scale bars correspond to 10 µm. The bottom panels show phase images for the same conditions. (**G**) Cumulative xenograft tumor volume in mice injected with control cells (n=6) or H661-p190A (n=7), as well H661-p190A with CRISPR/Cas9-mediated knockout *TJP2* using two different sgRNAs (n=11 each). Statistical testing was performed using pairwise Student’s *t* test as indicated by brackets. ****p<0.001; nsd: not significantly different. (**H**) Kaplan-Meier survival plot for mice the cohort of mice described in (**G**). Statistical testing was performed using pairwise log-rank test as indicated by brackets. **p<0.25; ****p<0.001; nsd: not statistically different. (**I**) Quantification of immunohistochemistry to detect Ki-67 positive cells in tumors 3 weeks after injection of control cells or H661-p190A cells w/wo *TJP2*-ko. Data are presented as mean ± SEM with n=5 mice and >600 nuclei scored per condition. Statistical testing was performed using pairwise Student’s *t* test as indicated by brackets. ****p<0.001; nsd: not significantly different.

E-cadherin is an established constituent of the canonical Hippo pathway, which was originally demonstrated using mammary carcinoma MDA-MB-231 cells ^33^. We subsequently determined that p190A induces expression of E-cadherin, which in turn is necessary for p190A to activate LATS kinases ^20^. Vice versa, in MDA-MB-231 cells, p190A is required for E-cadherin to promote LATS activation ^20^. Here, we further queried whether ZO-2 is necessary for E-cadherin to activate LATS kinases. Indeed, knockout of *TJP2* reduced LATS activation in MDA-MB-231 cells expressing E-cadherin (**Figure S5A and Figure S5B**). Thus, both p190A and ZO-2 are essential for E-cadherin to activate the canonical Hippo pathway.

Next, we tested a role for ZO-2 in p190A-mediated tumor suppression in a xenograft assay. We injected nude mice with control or H661-p190A cells, as well as H661-p190A cells with CRISPR/Cas9-mediated knockout of *TJP2* using two distinct sgRNAs. As demonstrated repeatedly, expression of p190A in H661 cells attenuated tumorigenesis followed by complete shrinkage of tumors (**Figure 4G**). In contrast, H661-p190A cells with knockout of *TJP2* formed tumors that were only slightly and insignificantly smaller than tumors from control cells (**Figure 4G**). The differences in survival between tumors resulting from inoculation of control and H661-p190A cells with *TJP2* knockout were equally marginal and not statistically significant (**Figure 4H**).

We performed a histological examination of tumors from a separate cohort where all mice were sacrificed three weeks after injection, again prior to the disappearance of tumors from mice injected with H661-p190A cells (**Figure S6A**). By IHC, we verified that p190A and ZO-2 were expressed as expected, i.e., that staining for p190A was absent from control cell tumors but present in H661-p190A cells with or without *TJP2*-ko (**Figure S6B**). Moreover, staining for ZO-2 was absent from tumors formed by H661-p190A cells with *TJP2*-ko, but present in tumors from the two other groups (**Figure S6C**). Finally, we determined that while H661-p190A cells were quiescent, cells expressing p190A with *TJP2*-ko exhibited similar intensity and frequency of Ki-67 staining as control cells (**Figure 4I and Figure S6D**). Thus, ZO-2 is obligatory for p190A-mediated tumor suppression.

### RasGAP and ZO-2 play essential roles in transcriptomic alterations elicited by p190A

In published studies, we have established that p190A exerts tumor suppressor capacities via modulation of gene transcription. We therefore queried whether RasGAP and ZO-2 might influence transcriptomic alterations elicited by expression of p190A in H661 cells. To this end, we analyzed transcriptomes from control cells, p190A(wt) or p190(Y2F) expressing cells, as well as H661-p190A(wt) cells with knockout of *TJP1* or *TJP2*. Given the results of our functional studies above, we compared gene expression in H661-p190A(wt) cells to each of the other conditions. The analysis revealed that gene expression in p190A(Y2F) cells is highly similar to that of control cells both in terms of number of genes up- or downmodulated, as well as the identity of genes (**Figure 5A, Figure 5B and Table S1**). Moreover, that in H661-p190A(wt) cells, *TJP2* modulates expression of significantly more genes than *TJP1* (**Figure 5A**). The shift along the PC2 axis in the PCA plot shown in **Figure 5B** suggests that expression of a subset of genes are affected similarly in control, H661-p190A(Y2F) and H661-p190A(wt) cells with *TJP2* knockout relative to cells expressing wild type p190A only. We therefore performed GSEA analysis to identify shared gene signatures. Of the 15 top pathways in this analysis, four pathways were conserved: Interferon alpha and gamma responses, MTORC1 signaling and EMT (**Figure 5C**). Putative roles for Interferon alpha and gamma responses, as well as MTORC1 signaling will be addressed elsewhere. Based on our prior work ^20^, the shared presence of an EMT signature is highly reassuring. Accordingly, waterfall plots for control, p190A(Y2F) and p190A(wt) cells with *TJP2* knockout relative to cells expressing wild type p190A only showed highly similar profiles (**Figure 5D**). The commonality extended to key individual EMT genes: *TWIST1*, *ZEB1*, CDH1 and CDH2 (**Figure 5E**). Taken together, the transcriptomic data are consistent with the functional studies in establishing essential roles for RasGAP and ZO-2 in tumor suppressor capacities of p190A.

**Figure 5:**
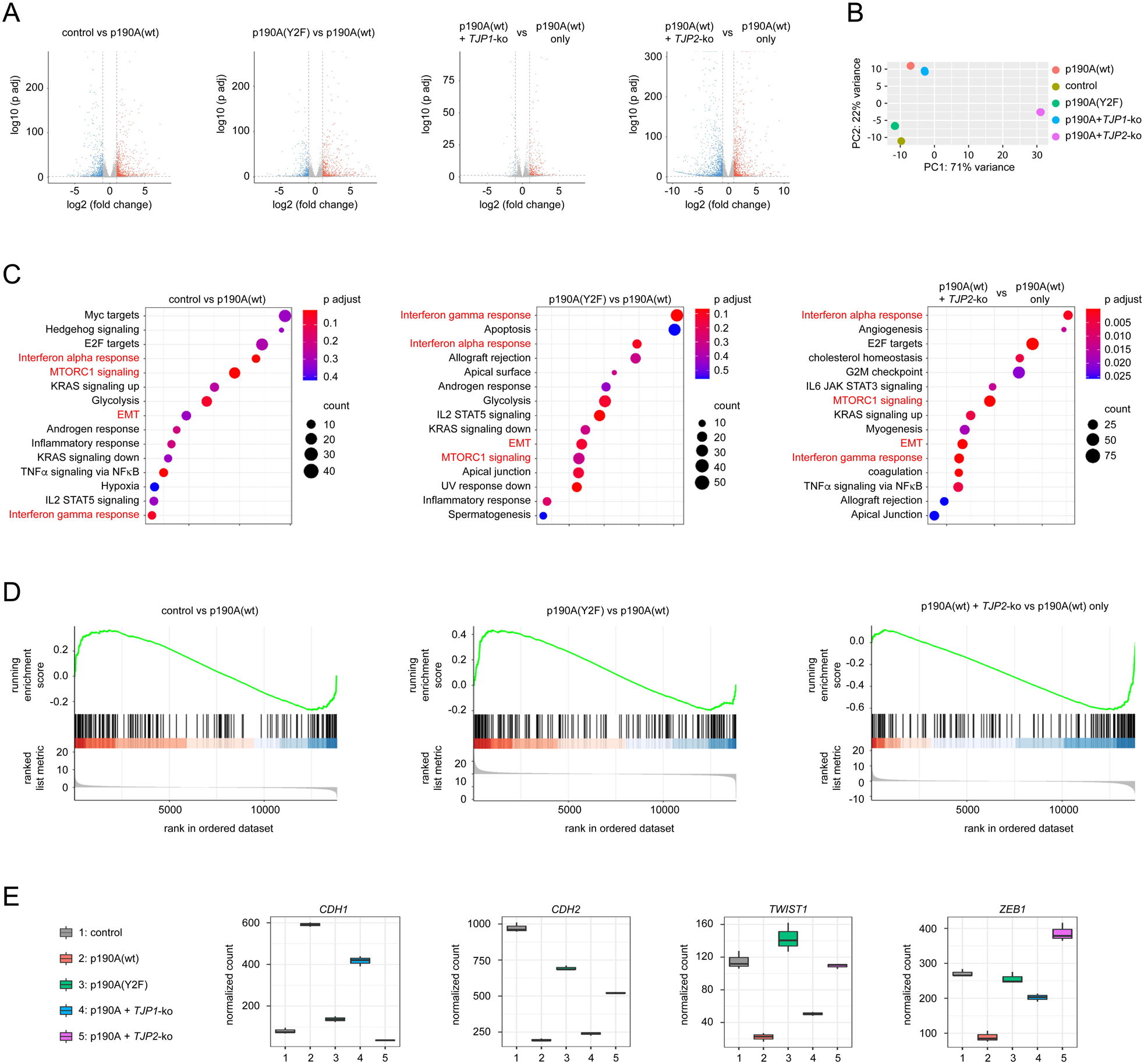
RasGAP and ZO-2 are required for transcriptomic alterations elicited by p190A. (**A**) Volcano plots of pairwise comparisons of differentially expressed genes as determined by RNA-seq in H661-p190A(wt) cells relative to control cells, or H661-p190A(wt) cells with or without knockout of *TJP1* or *TJP2*, as well as cells expressing p190A(Y2F). The threshold for differential expression was set at log2(fold change) >= 1 or <= -1with an adjusted p-value<0.05 (**B**) Principal component analysis (PCA) for RNA-seq gene expression in control H661 cells, as well as cells expressing p190A(wt) with or without *TJP1*-ko or *TJP2*-ko or cells expressing p190A(Y2F). Analysis was performed using DEseq2 on three biological replicates for each condition. (**C**) Dotplots showing the results of GSEA for 15 most enriched hallmark gene sets in pairwise comparisons of differentially expressed genes in H661-p190A(wt) cells relative to control cells, or H661-p190A(wt) cells relative to p190A(Y2F) cells, or H661-p190A(wt) cells with or without knockout of *TJP2*. (**D**) Waterfall plots of GSEA EMT gene sets in H661-p190A(wt) relative to control cells, H661-p190A(wt) relative to p190A(Y2F) cells, or H661-p190A(wt) cells with or without knockout of *TJP2*; p value = 0.044, 0.016, and < 0.001, respectively. (**E**) Expression of the key EMT genes, *CDH1*, *CDH2*, *TWIST1* and *ZEB1*, as determined by RNA-seq for the following conditions: control, H661-p190A(wt), and p190A(Y2F), as well as H661-p190A(wt) with *TJP1* or *TJP2* knockout.

### Expression of *TJP1/2* differentiates survival of patients with *ARHGAP35* alteration in LUAD

We mined TCGA data for putative links between expression of *ARHGAP35* and *RASA1*, as well as *TJP1* and *TJP2*. Given that our functional studies are conducted with NSCLC cells, we focused on lung adenocarcinoma (LUAD) where *ARHGAP35* is frequently altered, especially in oncogene negative tumor samples ^34^. In gsea-msigdb.org, there is a highly significant correlation between *ARHGAP35* expression in LUAD samples and Hippo signaling (**Figure 6A**). This observation provides crucial validation in human tumor samples of links between *ARHGAP35* and the canonical Hippo pathway that we previously have established using cultured cells ^19,20^. We then examined expression of *ARHGAP35* and *RASA1*, as well as *TJP1* and *TJP2* in tumor samples and uninvolved normal tissue from the same patients. Expression of *ARHGAP35*, *TJP1* and *TJP2* were all significantly downmodulated in tumor samples relative to matched normal tissue (**Figure 6B and Figure S7A**). In contrast, *RASA1* expression was significantly increased in tumor tissue (**Figure 6B**). Reasons for the latter observation are not clear, but could be linked to the established role for RasGAP in angiogenesis ^35^. Next, we stratified patients based on *TJP1/2* transcript levels in tumors and queried the impact of *ARHGAP35* expression on survival in the respective patient populations. Strikingly, low *ARHGAP35* expression correlates with shorter survival in patients with high but not low transcript levels of *TJP1/2*. This discovery is consistent with the requirement for ZO-2 in tumor suppressor signaling elicited by p190A. The apparent role for ZO-1 might be due to cell type-dependent differences in the respective roles for ZO-1 and ZO-2 for p190A function. Alternatively, it could also be explained by strong positive correlation between *TJP1* and *TJP2* expression in the TCGA LUAD dataset (**Figure S7B**).

**Figure 6:**
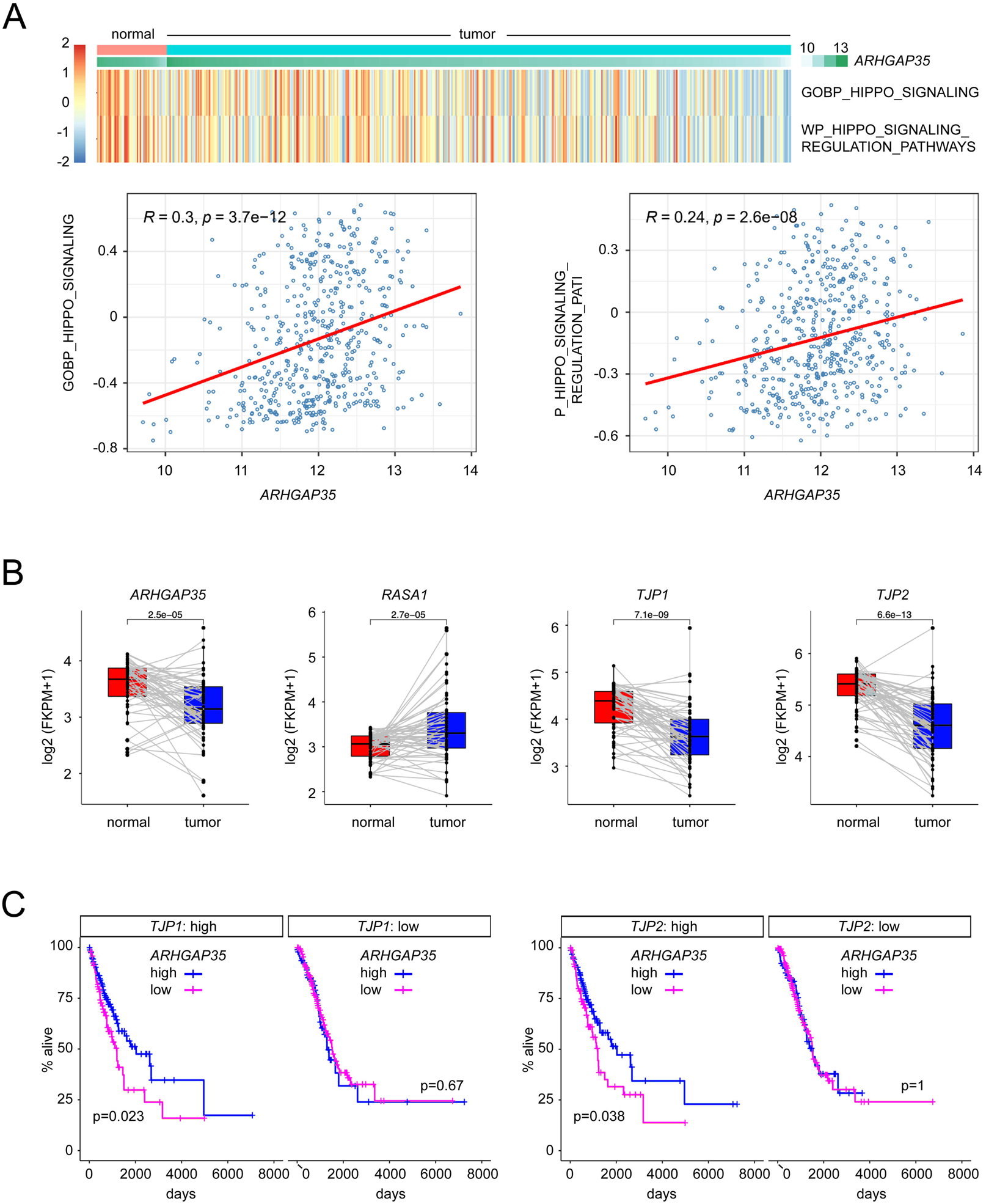
Low *ARHGAP35* expression is associated with reduced lifespan in patients with high, but not low, levels of *TJP1/2* transcripts. (**A**) Upper panel: Heat map of activity scores of Hippo pathways in 524 LUAD and 59 normal solid tissue samples retrieved from TCGA. Raw counts data were used for analyzing Hippo pathway activity using the GOBP_HIPPO_SIGNALING and WP_HIPPO_SIGNALING_ REGULATION_PATHWAYS gene sets from MsigDB by GSVA. Lower panel: Correlation between *ARHGAP35* expression (VST transformed data by DESeq2) and Hippo signaling in 524 LUAD solid tissue samples. (**B**) Differences in *ARHGAP35*, *RASA1*, *TJP1*, and *TJP2* transcript levels in matched normal and tumor tissue from patients with LUAD. (**C**) Survival of patients with high and low *ARHGAP35* expression stratified according to high versus low transcript levels of *TJP1* or *TJP2*. Survival data from a total of 511 patients were available for this analysis. Medium value was used as cutoff for separating *ARHGAP35, TJP1* or *TJP2* high and low expression groups.

## Discussion

*ARHGAP35* encoding p190A remains very sparingly studied in the context of cancer despite being highly altered by both deletion and mutation ^1–3^. The spectrum of alterations is suggestive of a tumor suppressor function. Accordingly, our past studies have determined that p190A is indeed a tumor suppressor and, consistently with this role, an upstream activator of the Hippo pathway ^19,20^. Here, we aimed to identify a physical link between p190A and established constituents of the Hippo pathway. To this end, we performed a mass spec analysis to identify p190A interacting proteins in H661 cells, where we have shown p190A activates LATS kinases. This analysis revealed RasGAP and Zonula Occludens proteins ZO-1/2 to rank among the most frequently interacting proteins.

A cDNA encoding p190A was originally cloned via the interaction with RasGAP, hence validating the results of our mass spec analysis ^24^. In the present study, we demonstrate that RasGAP is necessary for p190A to activate LATS kinases, elicit MET, induce CIP and suppress tumorigenesis. This discovery is important, because it expands the role of RasGAP as tumor suppressor beyond promoting hydrolysis of GTP-bound Ras proteins. Thus, the p190A-RasGAP complex may serve as a node for coordinating Hippo and Ras signaling. To this end, Campbell et al. strikingly demonstrated that *ARHGAP35*, *RASA1* and *LATS1* rank among the most frequently altered genes in oncogene-negative LUAD ^34^. Our present study establishes that p190A, RasGAP and LATS1 signal in the same pathway. By inference, loss of function alterations in this pathway may drive oncogene-negative LUAD by activating YAP/TAZ transcriptional co-activators, which also have been shown to confer oncogenic capacities in Ras-resistant cancer ^37,38^.

Next, we demonstrate that RasGAP is necessary for the novel interaction of p190A with the Zonula Occludens proteins ZO-1 and ZO-2 that serve as transcriptional modulators of ZONAB and TEAD transcription factors ^39,40^. ZO-2 moreover represents an integral constituent of the Hippo pathway via its direct interaction with YAP/TAZ and potentially LATS kinases ^41–43^. Using MDCK cells expressing GFP-tagged ZO-1 and derivatives thereof, we demonstrate that p190A interacts with C-terminal region of ZO-1 harboring the GuK and ABR motifs, i.e. distant from the PDZ1 domain in the N-terminus that binds YAP/TAZ ^41,42^. Moreover, we show that ZO-1 and ZO-2 recruit p190A to lateral membranes of MDCK cells. We have hitherto been unable to determine the subcellular localization of these constituents in the cancer cell lines studies herein. However, the observations from MDCK cells establish that ZO-1 and ZO-2 possess the capacity to dictate the subcellular localization of p190A to a site relevant to CIP. Functionally, our data demonstrate that ZO-2 is essential for p190A to activate LATS kinases, elicit MET, induce CIP and suppress tumorigenesis. ZO-2 is also required for E-cadherin to promote LATS activation, which we previously have shown to require p190A ^20^. Moreover, ZO-2 is necessary for the transcriptional response associated with tumor suppression elicited by expression of p190A. Finally, we determine that in LUAD samples where *ARHGAP35* expression correlates positively with Hippo signaling, low expression of *ARHGAP35* is associated with worse prognosis, but only in patients with high *TJP2* expression in tumor tissue, consistent with *ARHGAP35* acting as a *TJP2*-dependent tumor suppressor gene in humans.

Work from other laboratories have established that Rho-ROCK-mediated actomyosin contraction represses activation of LATS kinases ^44–46^, a mechanism that directly contributes to driving human malignancies ^47^. This pathway, also referred to as mechano-transduction, attenuates direct binding between Nf2/Merlin and LATS kinases to promote YAP-mediated gene transcription ^48^. Thus, inhibition of Rho signaling is essential to activate LATS kinases to promote growth control. Experimentally, this is typically achieved with global inhibitors of actin polymerization such as C3 toxin or latrunculin B. However, in physiological settings, Rho signaling must be fine-tuned to coordinate a vast array of processes that requires simultaneous activation and inactivation of Rho activity at distinct subcellular sites ^49^. Our past work has made p190A strong candidate for a key enzyme to repress mechano-transduction in activation of LATS kinase ^19^. Yet, hitherto, we have been unable to identify physical association between p190A and established constituents of the Hippo pathway. However, as summarized in the graphical abstract accompanying this article, our present demonstration of an interaction between the RasGAP-p190A complex and ZO-2 provides this missing link, such that the RasGAP-p190A complex can now be considered a *bona fide* constituent of the Hippo pathway. This is further supported by the demonstration by Cui et al of a direct interaction between RasGAP and Nf2/Merlin ^50^, although its significance to Hippo signaling remains to be determined.

In summary, this work elucidates a novel mechanism whereby the p190A-p120 complex via interaction with ZO-2 exerts tumor suppressor capacities. From a rather obscure role in cancer, p190A is emerging as pivotal modulator of the Hippo pathway with physical and functional links to established constituents of cell-cell junctions that are known to modulate the activity of LATS kinases and YAP/TAZ transcriptional co-activators ^51^.

## Supplemental Information

Supplemental Information includes experimental methods, seven figures and one table.

## Acknowledgements

We are grateful to Jerrold R. Turner for providing MDCK cell lines, Phi Luong for critical reading of the manuscript, and to Suzanne L. White and Dr. Lay-Hong Ang, BIDMC confocal imaging and IHC core for assistance with histology procedures. This work utilized Harvard Digestive Diseases Center (HDDC) core facilities (NIH P30DK034854) and was funded by NIH R01 CA205158, as well as an endowment from the Roy and Lynne Frank Foundation (to Steen H. Hansen).

