## Supplemental information for "p120 RasGAP and ZO-2 are essential for Hippo signaling and tumor suppressor function mediated by p190A RhoGAP"

**Methods**

**Reagents**
The following reagents were used for this work were the following: Blasticidin (EMD Millipore); Calyculin A (Cell Signaling Technologies); DMEM (Corning); DMEM/F12 1:1 (Gibco); DRAQ5 (Biostatus); fetal bovine serum (FBS; Atlanta Biologicals); Luna Universal qPCR Master Mix (New England Biolabs); fish skin gelatin ( Sigma-Aldrich); FluorSave™ (EMD Millipore); Fugene6 (Promega); goat serum (Gibco); Omnifect (Transomic technologies); PageRuler™ prestained protein ladder 10-180 kD (Thermo Fisher); puromycin (Sigma-Aldrich); phalloidin/Alexa$^{488}$ and phalloidin/Alexa$^{594}$ (Invitrogen); Polybrene (Santa Cruz Biotechnology); Puromycin (Sigma Aldrich); Trypsin-EDTA, 0.25% (Gibco).

**Antibodies**
Antibodies used for this study were as follows:

| Antigen | Species | Company | Cat. no. |
|---|---|---|---|
| Cyclin A | rabbit polyclonal | Santa Cruz Biotechnologies | sc-751 |
| E-Cadherin | mouse monoclonal | BD Biosciences | 610182 |
| ERK1 | mouse monoclonal | Santa Cruz Biotechnologies | sc-271269 |
| ERK1/2 | rabbit polyclonal | Sigma Aldrich | M5670 |
| pERK1/2 | rabbit monoclonal | Cell Signaling Technologies | 4370S |
| GFP | mouse monoclonal | Santa Cruz Biotechnologies | sc-9996 |
| Ki-67 | rabbit polyclonal | Thermo Fisher Scientific | RM9106S |
| pLATS1(S909) | rabbit monoclonal | Cell Signaling Technologies | 9157S |
| N-cadherin | mouse monoclonal | Cell Signaling Technologies | 14215S |
| MEK | rabbit monoclonal | Cell Signaling Technologies | 9126S |
| pMEK | rabbit monoclonal | Cell Signaling Technologies | 9128S |
| p190A | mouse monoclonal | BD Biosciences | 610150 |
| p190B | mouse monoclonal | BD Biosciences | 611613 |
| ZO-1 | rabbit monoclonal | Cell Signaling Technologies | 8193S |

| | | | |
|---|---|---|---|
| ZO-1 | rabbit monoclonal | Cell Signaling Technologies | 13663S |
| ZO-1 | rabbit polyclonal | Proteintech | 21773-1-AP |
| ZO-2 | rabbit monoclonal | Cell Signaling Technologies | 2847S |
| ZO-2 | rabbit polyclonal | Proteintech | 18900-1-AP |
| ZO-2 | rabbit polyclonal | Boster | PA1957 |

Secondary antibodies obtained from Invitrogen were the following: goat anti-mouse/Alexa$^{555}$; goat anti-rabbit/Alexa$^{488}$; goat anti-mouse/HRP; goat anti-rabbit/HRP.

**Plasmid constructs**

wild type p190A and p190A(Y2F) expression constructs were synthesized with an N-terminal Myc-tag by Genewiz and cloned into the lentiviral vector pUltra-hot from Addgene, plasmid #24130. The entire cDNAs for wild-type or p190A(Y2F) forms were verified by Sanger sequencing. Lentiviral pZIP vectors encoding shRNAs targeting human p190A were purchased from transOMIC technologies Inc; cat. no. TRHS1000-35 (*ARHGAP35*/p190A) and validated as previously described [39]. For CRISPR/Cas9-mediated knockout of *RASA1*, *TJP1* and *TJP2*, we utilized the following guide sequences:

*RASA1* encoding p120 RasGAP
sgRNA#1 TCC CGT GTC GGG TGA AGA TAC CCG
sgRNA #2 TCC CTC TGG ATG GAC CAG AAT ACG

*TJP1* encoding ZO-1
sgRNA #1 TCC CGG AAA ATG ACC GAG TTG CAA
sgRNA #2 TCC CTG ACC GCC TGT CTG ACC GCG

*TJP2* encoding ZO-2
sgRNA #1 TCC CAC GGG TCT GGC AAC TAA AGA
sgRNA #2 ACC GAA AGA TGG CAA CCT TCA CGA

sgRNA#2 for *TJP2* in pLentiGuide-Puro was purchased from Addgene (#77828). All other sgRNAs were cloned into FgH1tUTG (Addgene #70183), which expresses EGFP for selection by FACS.

**Cell culture, transfection, transduction, and selection**

NCI-H661 and NCI-H226 cell lines were propagated in DMEM/F12 1:1 supplemented with 10% FBS. 293T, MDA-MB-231 and MDCK cells grown in DMEM with 10% FCS. Transfections were performed using Omnifect or Fugene6 according to the manufacturer's instructions. For lentiviral transduction, one 100-mm dish with 70% confluent 239T cells was transfected with 2 µg each of VSV-G and PAX2 encoding plasmids, as well as 2-ug transfer vector. The medium was replaced after 24 hours, and medium containing lentiviral particles harvested after 48 hours. Following filtration through a 0.45-µm filter, the medium was supplemented with 8 µg/ml polybrene and added to target cells. Transduced cells were enriched either by FACS sorting for EGFP or mCherry expression, or by drug selection with 10 µg/ml blasticidin or 2 µg/ml puromycin for 10 days. Selected cells were expanded to generate frozen stock and then used for experimentation.

**Confocal microscopy**

Samples were washed once with PBS and fixed in 3.7% formalin containing 1% methanol for 10 min at room temperature. After washing three times with PBS, samples were incubated 30 min in PBS containing 10% normal goat serum, 0.2% fish skin gelatin and 0.1% Triton X-100 (PBS-NGS). Next, samples were incubated with primary antibodies diluted in PBS-NGS for 1h, rinsed extensively with PBS containing 0.1% Triton X-100 for 30 mins, and then incubated with secondary antibodies diluted in PBS-NGS for 40 mins. Following additional extensive rinsing for 30 mins, samples were stained for 15 mins with DRAQ5 diluted 1:300 in PBS to detect nuclei and, when relevant, phalloidin/Alexa$^{488}$ or phalloidin/Alexa$^{594}$ to visualize polymerized actin. After final rinsing in PBS, samples were mounted with coverslips using FluorSave.

**Tumorigenesis in nude mice**

6-7 weeks old female outbred homozygous nude Foxn1$^{nu}$/Foxn1$^{nu}$ were obtained from The Jackson Laboratory Cat. no. 007850) and acclimatized for 2-3 days. Mice were then ear-tagged and injected with 100µl Matrigel containing 5x10$^{6}$ control or H661-p190A cells in the right flank. Mice were followed for up to 25 weeks after injection, during which weight and period tumor size (length x width) were measured 1-2 times per week and the condition of mice inspected. When tumors reached a maximum length of ≥8-mm, mice were euthanized by $CO_2$ asphyxiation per institutional guidelines and tumors were removed for histology. These procedures were conducted according to BCH IACUC-approved protocol #3319. In parallel, smaller cohorts were established in which all mice, irrespective of tumor size were euthanized three weeks after injection of tumor cells.

**Histology**

Tumors were removed from euthanized mice and fixed in 10% formalin overnight, dehydrated and embedded in Tissue Prep 2 paraffin. Samples were sectioned on a rotary microtome, and H&E staining was performed in an Autostainer. Retrieval of antigens was performed by boiling slides for 10 minutes in 10mM Sodium Citrate buffer in a pressure cooker. Sections were blocked with 5% normal donkey serum (Jackson ImmunoResearch Lab Inc, West Grove PA) for one hour at room temperature. To quantify Ki-67, sections were incubated with rabbit anti-Ki-67 antibody overnight at 4$^{o}$C. Sections were then washed in TBS/TBST and incubated with Alexa$^{488}$-conjugated donkey anti-rabbit secondary antibody diluted 1:300. Next, sections were counter-stained with Hoechst 33342, washed with TBS/TBST and mounted in Prolong Gold anti-fade mounting media (Invitrogen). Finally, Cell Profiler (Broad Institute) was used to define and count Ki-67 and Hoechst 33342 positive nuclei, respectively. Immunohistochemistry to detect p190A, ZO-2 and Ki-67 was performed using similar methods, except that incubation with primary antibodies was followed by HRP-conjugated goat anti-rabbit/mouse secondary antibody followed by staining with hematoxylin. H&E staining was performed using standard methods.

**Real-time qPCR**

Total RNA was extracted from cell pellets using Qiagen RNeasy Mini and QIAshredder kits according to the manufacturer's instructions. Next, cDNA was synthetized using Bio-Rad iScript cDNA Synthesis Kit according to the manufacturer's instructions. qRT-PCR was then performed with the One Step plus Sequence Detection System using Fast SYBR green master mix reagent. Finally, gene expression levels were normalized to the "housekeeping" genes *HPRT1* and *RPS18*. Primer sequences were as follows:

| Gene | Forward primer (5′-3′) | Reverse primer (5′-3′) |
|---|---|---|
| *CDH1* | GTCACTGACACCAACGATAATCCT | TTTCAGTGTGGTGATTACGACGTTA |
| *CDH2* | CCTCCAGAGTTTACTGCCATGAC | GTAGGATCTCCGCCACTGATTC |
| *CTGF* | GAAGCTGACCTGGAAGAGAACA | CGTCGGTACATACTCCACAGAA |
| *CYR61* | GAGTGGGTCTGTGACGAGGAT | GGTTGTATAGGATGCGAGGCT |
| *TWIST1* | GCCAGGTACATCGACTTCCTCT | TCCATCCTCCAGACCGAGAAGG |
| *ZEB1* | GGCATACACCTACTCAACTACGG | TGGGCGGTGTAGAATCAGAGTC |
| *SNAIL2* | ATCTGCGGCAAGGCGTTTTCCA | ATCTGCGGCAAGGCGTTTTCCA |
| *HPRT1* | TTGCTTTCCTTGGTCAGGCA | ATCCAACACTTCGTGGGGTC |
| *RPS18* | CTTTGCCATCACTGCCATTAAG | TCCATCCTTTACATCCTTCTGTC |

**Genome-wide mRNA expression profiling**

Total RNA was isolated from $1x10^6$ NCI-H661 cells per sample using the RNeasy kit (Qiagen). Sample integrity was verified on an Agilent Technologies 2100 Bioanalyzer using the Agilent RNA 6000 Nano kit with an RNA integrity number (RIN) above 9.5 as threshold for acceptance. Next, samples were shipped to BGI Genomics Co. Ltd for further processing. In brief, total RNA was subjected to Oligo dT selection for enrichment of mRNA followed by reverse transcription and second strand synthesis. Following cDNA synthesis and library preparation, 50 bp end sequencing was performed on the BGISEQ-500 platform with a minimum of 20M clean reads per sample. Sequence reads were then filtered with SOAPnuke software to remove reads containing adaptors or unknown bases, as well as low-quality reads. Next , genome mapping of filtered reads to reference genome GRCh38 was performed using HISAT2 and Bowtie2 software [65,66]. Gene expression levels were calculated with RSEM [67]. Finally, differential gene expression was detected with NOIseq [68].

**RNA-seq data analysis and TCGA data analysis**

Analyses and figure generation were conducted within the RStudio using R version 3.63. Raw data of RNA-seq were normalized and processed using R package DEseq2 XX. Principal Components Analysis (PCA) was plot using plotPCA function. Differentially expressed genes (DEGs) were analyzed using DESeq2. Genes with baseman > 5, false discovery rate (FDR) < 0.05 and log2 (fold change) value >= 1 were considered significantly regulated. Hallmark gene sets were downloaded from MSigDB and Gene set enrichment analysis (GSEA) was performed with Clusterprofiler package in R to determine the enrichment of pathways. The data are deposited in GEO submission GSE212619.

For GDC TCGA Lung Adenocarcinoma (LUAD) cohort, both raw counts and processed FPKM-UQ data of were downloaded from UCSC Xena Browser Datasets. RNA-seq data and matched clinical data were available for 524 primary tumor and 59 adjacent normal solid samples. FPKM-UQ normalized data were used for comparing the gene expression levels between tumor and normal tissues. Raw counts data were used as input and processed using GSVA package for analyzing Hippo pathway activity. Hippo pathway gene sets (GOBP_HIPPO_SIGNALING and WP_HIPPO_SIGNALING_REGULATION _PATHWAYS) were downloaded from MSigDB. Survival data were available for 511 patients of this cohort and was used for survival analysis.

**Data and Code Availability**

The RNA-seq data have been deposited in NCBI's Gene Expression Omnibus and can be accessed using GEO Series accession number GSE212619.

**Statistical analyses**

Student t tests (unpaired, two-tailed, unequal variance) and log-rank tests were performed as described previously [69]. In all figures *, **, ***, and **** denote $p<0.05$, 0.025, 0.01, and 0.001 respectively.

**Figure S1**

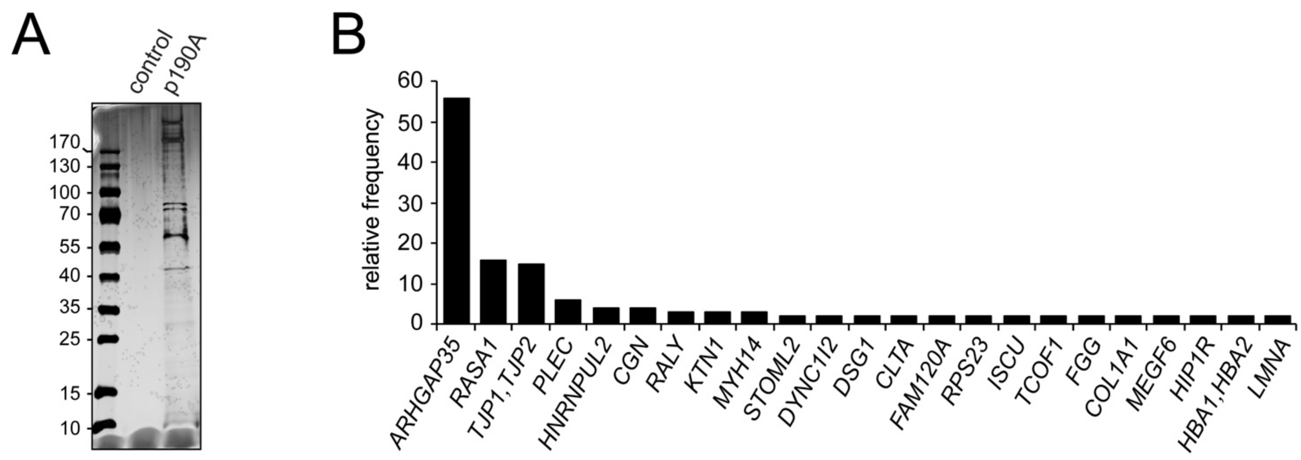


C

*ARHGAP35*: p190A RhoGAP
*RASA1*: p120 RasGAP
*TJP1,TJP2*: Zonula Occludens 1,
*PLEC*: Plectin
*HNRNPUL2*: Heterogeneous Nuclear Ribonuclear Protein U Like 2
*CGN*: Cingulin
*RALY*: RALY Heterogeneous Nuclear Ribonuclear Protei
*KTN1*: Kinectin 1
*MYH14*: Myosin Heavy Chain 14
*STOML2*: Stomatin Like 2
*DYNC1I2*: Dynein Cytoplasmic 1 intermediate Chain 2
*DSG1*: Desmoglein 1
*CLTA*: Clathrin Light Chain A
*FAM120A*: Family With Sequence Similarity 120A
*RPS23*: Ribosomal Protein S23
*ISCU*: Iron Sulfur Cluster Assembly Enzyme
*TCOF1*: Treacle Ribosome Biogenesis Factor 1
*FGG*: Fibrinogen Gamma Chain
*COL1A1*: Collagen Type I Alpha 1 Chain
*MEGF6*: Multiple EGF Like Domains 6
*HIP1R*: Huntingtin Interacting Protein 1 Related
*HBA1,HBA2*: Hemoglobin Subunit Alpha 1,2
*LMNA*: Lamin A/C

**Figure S1: Mass spectrometry analysis of p190A-interacting proteins in H661 cells.** (**A**) Silver stain of SDS-PAGE gel following electrophoresis of 9E10 anti-Myc-Agarose immunoprecipitated proteins from either control cells harboring defined *ARHGAP35* alterations resulting in complete loss of p190A, or from H661 cells reconstituted with Myc-tagged p190A (H661-p190A). (**B**) Relative frequency of peptides detected by mass spec from H661-p190A cells after elimination of peptides detected in control cells. (**C**) Full names of proteins from gene abbreviations in panel (**B**).

**Figure S2**

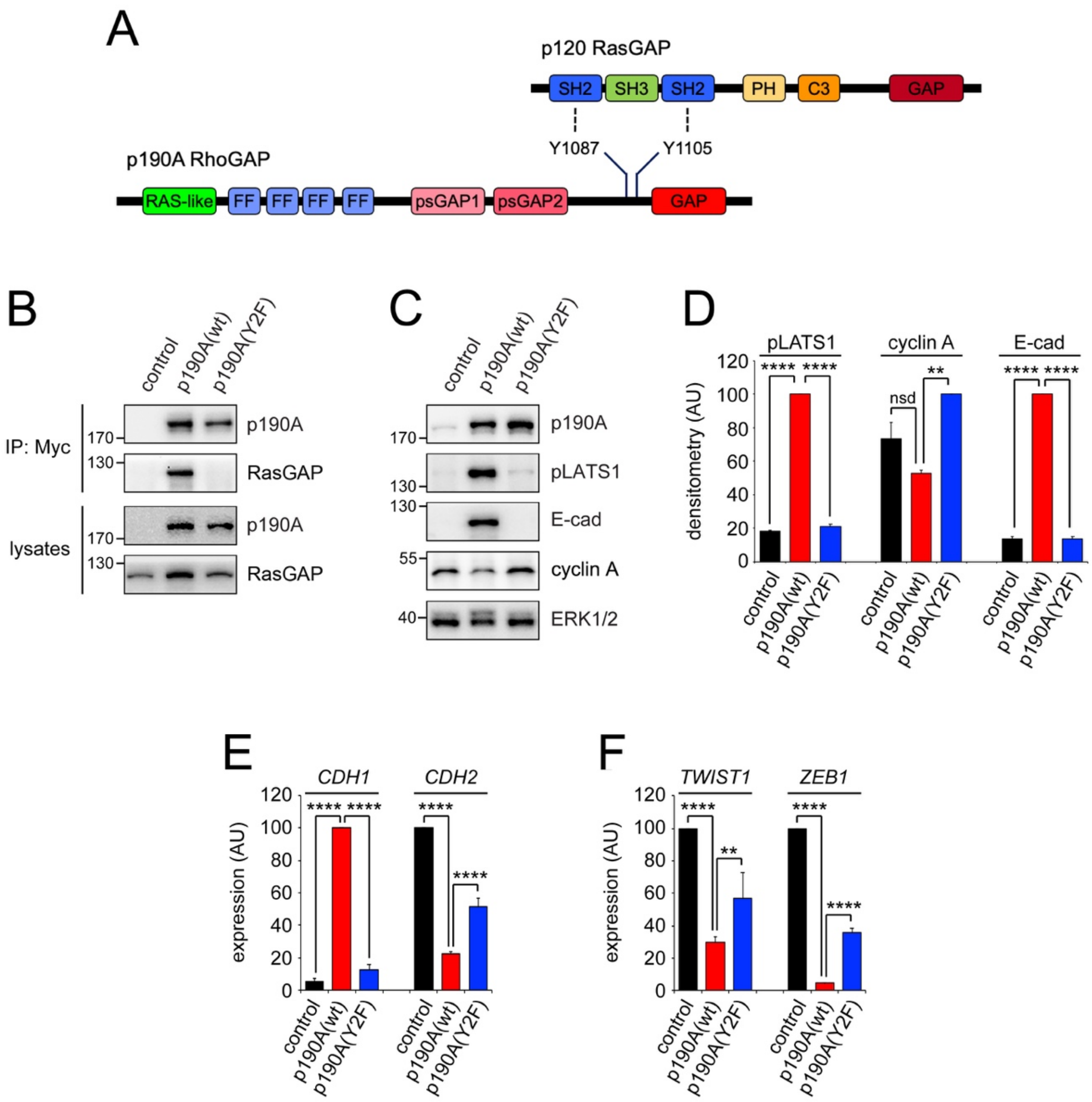


**Figure S2: Interaction of p190A with RasGAP is necessary for p190A to activate of LATS1 kinase, promote CIP and elicit MET in H226 cells.** (**A**) Cartoon illustrating domains and motifs mediating direct interaction between p190A and RasGAP. (**B**) Western blots of co-IPs and corresponding whole cell lysates to detect p190A and RasGAP, pLATS1 in control cells, as well as H226 cells reconstituted with either p190A(wt) or p190A(Y2F) defective in RasGAP-binding. (**C**) Western blots of whole cell lysates to detect p190A, pLATS1, cyclin A, E-cadherin and ERK1/2 in control cells, as well as H226 cells reconstituted with either p190A(wt) or p190A(Y2F). (**D**) Quantification by densitometry of pLATS1, cyclin A and E-cadherin levels detected by western blotting as shown in (**C**). Data are presented as mean ± SD (n=3). (**E**) Transcript levels for *CDH1* and *CDH2* in H226 cells as determined by qPCR. Data are presented as mean ± SD (n=3). (**F**) Transcript levels for *TWIST1* and *ZEB1* as determined by qPCR. Data are presented as mean ± SD (n=3). All statistical testing for data presented in this figure was performed using pairwise Student's *t* test as indicated by brackets. **$p<0.025$; ****$p<0.001$; nsd, not significantly different.

**Figure S3**

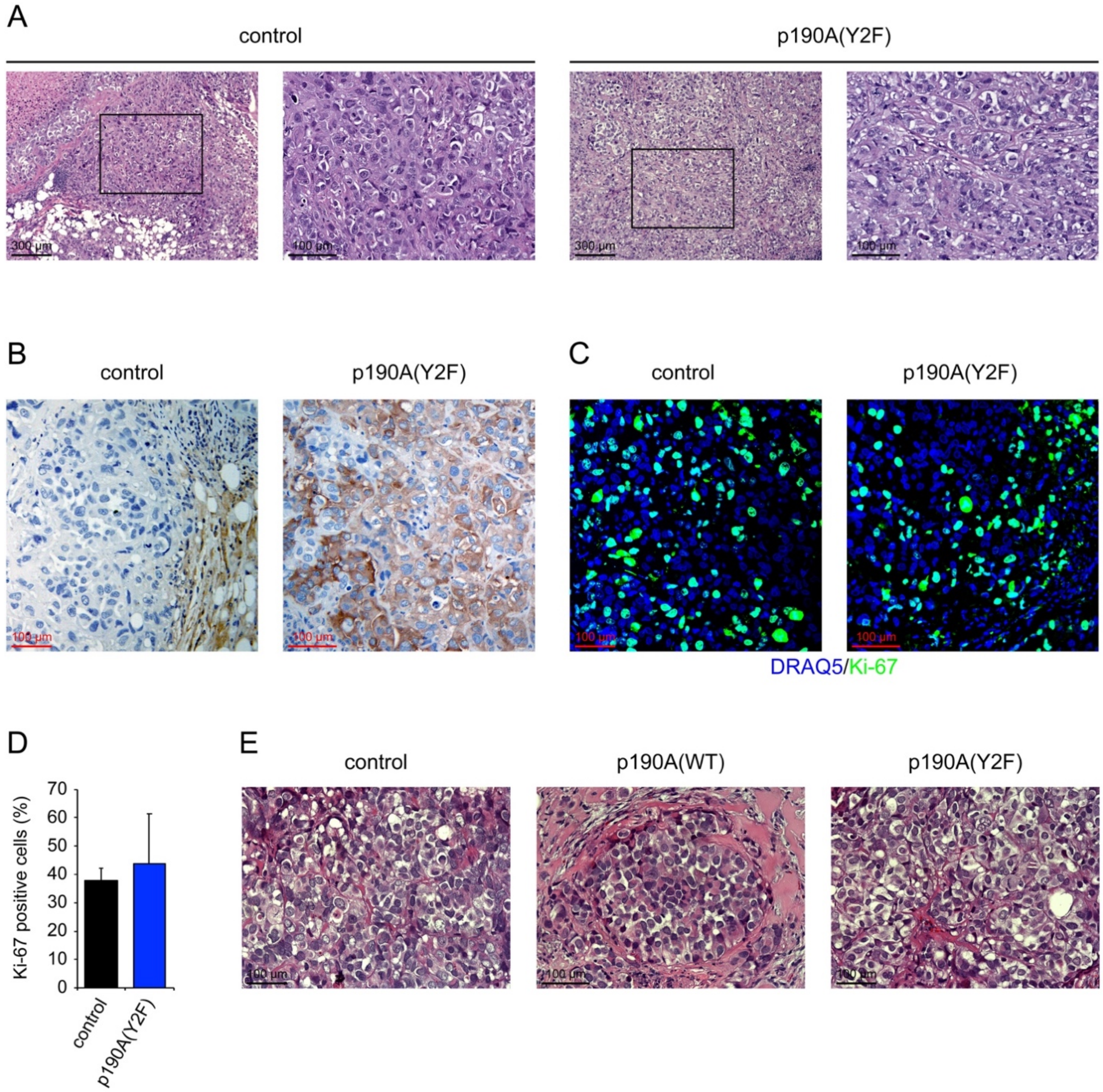


**Figure S3: H&E staining and immunohistochemistry to detect p190A and Ki-67 antigen in tumors from mice inoculated with control, H661-p190A(wt) and H661-p190A(Y2F) cells.** (**A-D**) Analyses of tumors harvested from mice from mice injected with control or H661-p190A(Y2F) from the cohort for which tumor volume and Kaplan-Meier survival plot are shown in main **Figure 2A and Figure 2B**, respectively. (**A**) H&E staining of control and p190A(Y2F) tumors as indicated. (**B**) Immunohistochemistry to detect p190A protein in control and p190A(Y2F) tumors. (**C**) Immunofluorescence to detect Ki-67 antigen in control and p190A(Y2F) tumors. (**D**) Quantification of Ki-67 staining from three tumor each from control and p190A(Y2F) tumors. There is no significant difference in the number of Ki-67 positive nuclei, as determined by Student's *t* test. (**E**) H&E staining from the 3-week cohort of mice for which p190A and Ki-67 immunohistochemistry analyses are shown in main **Figures 2C-F**.

**Figure S4**

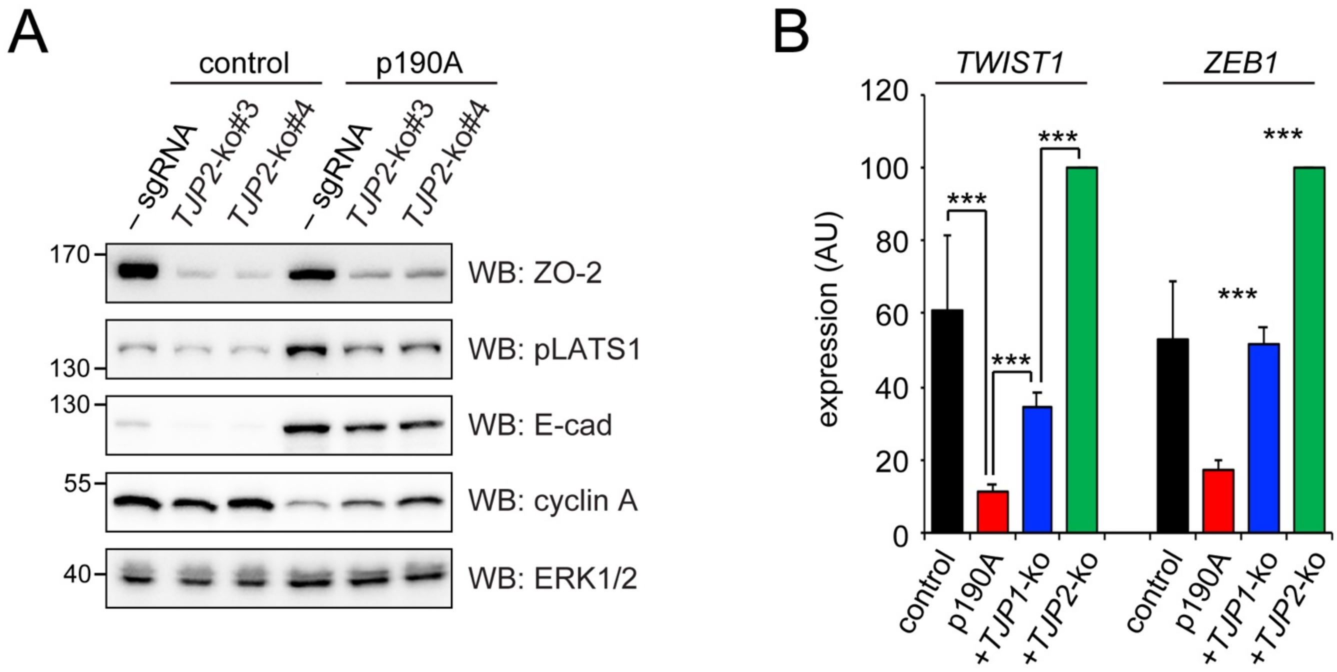


**Figure S4: ZO-2 is required for p190A activate LATS kinases, promote CIP and elicit EMT.** (**A**) Western blots to detect ZO-2, pLATS1, E-cad, cyclin A and ERK1/2 in control and H661-p190A cells with or without *TJP2* knockout using two additional distinct sgRNAs from those shown in Figure 4. (**B**) qPCR to detect expression of TWIST1 and ZEB1 genes in control cell and H661-p190A cells with or without knockout of *TJP1* or *TJP2*.

**Figure S5**

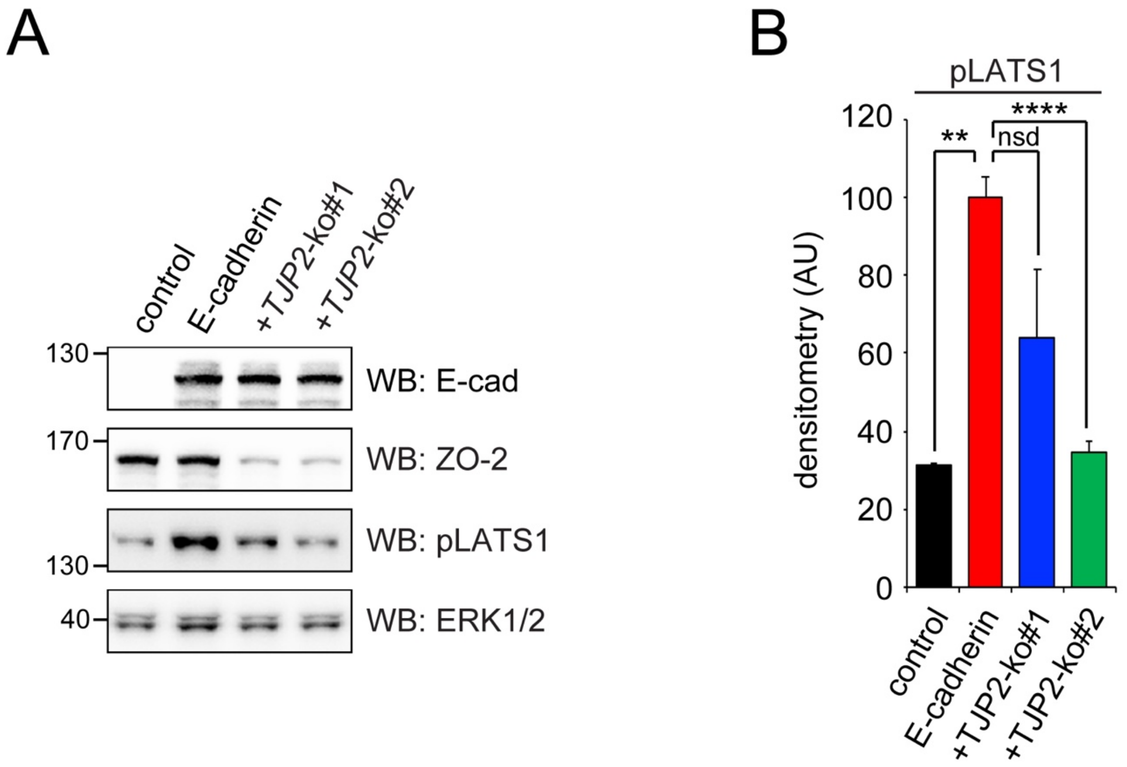


**Figure S5: ZO-2 is necessary for E-cadherin to activate LATS1 in MDA-MB-231 cells.** (**A**) Western blots to detect E-cad, ZO-2, pLATS1 and ERK1/2 in MDA-MB231 cells with or without constitutive expression of exogenous E-cadherin and with or without knockout of endogenous *TJP2*. (**B**) Quantification by densitometry of E-cadherin, and N-cadherin levels detected by western blotting as shown in (**A**).

**Figure S6**

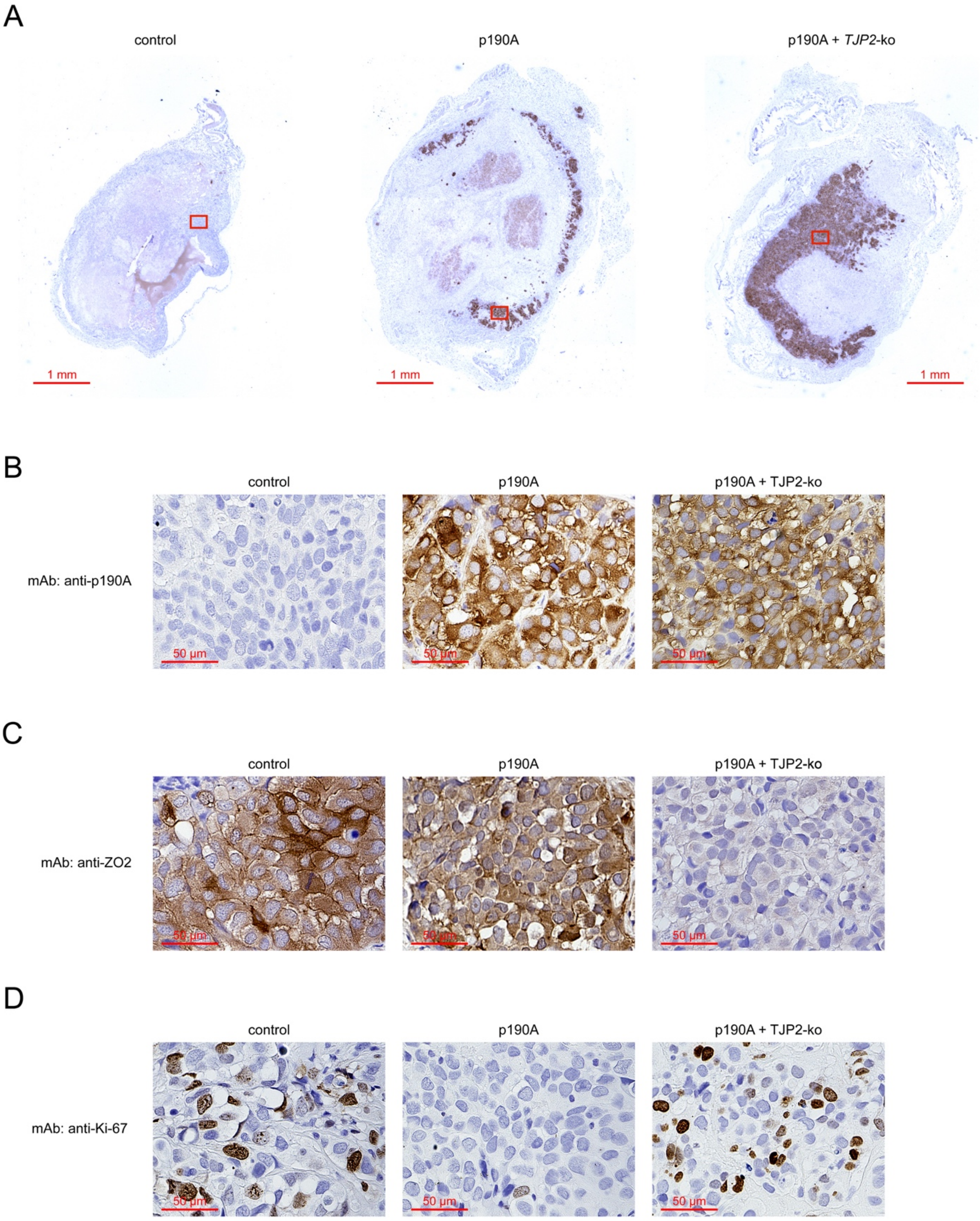


**Figure S6: ZO-2 is required for the tumor suppressor function of p190A.** (**A**) Immunohistochemistry to detect p190A in tumors 3 weeks after injection of control cells or cells expressing wild type p190A with or without knockout of *TJP2*. (**B**) 20x magnifications of the areas contained within the red boxes in (**A**). (**C**) Immunohistochemistry to detect ZO-2. (**D**) Immunohistochemistry to detect Ki-67.

**Figure S7**

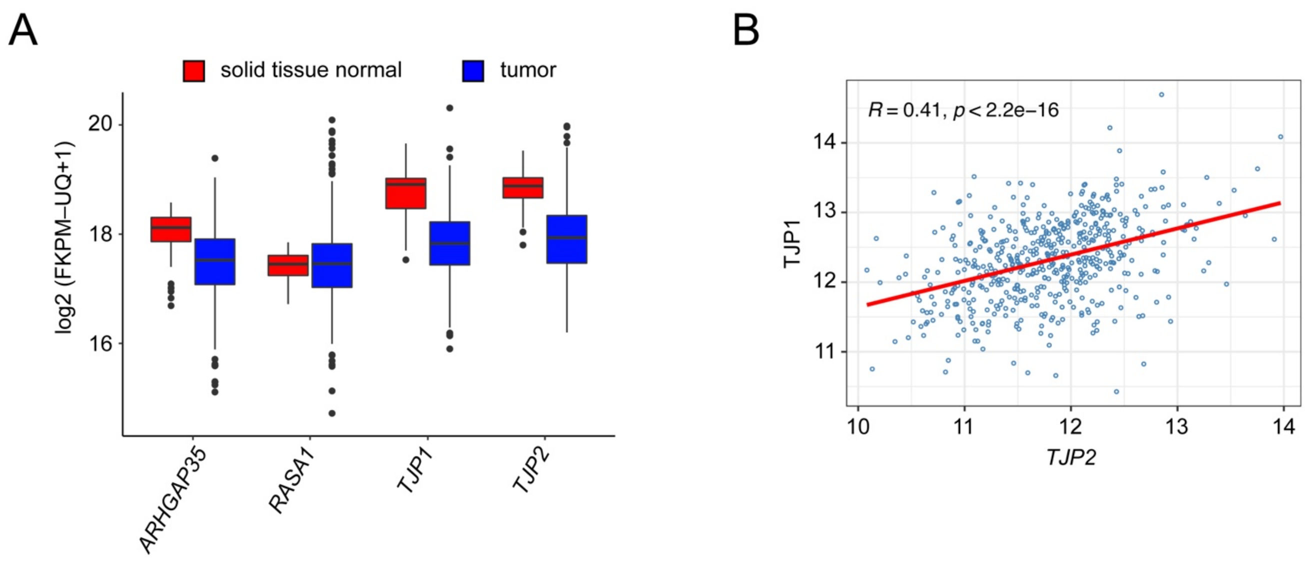


**Figure S7: Expression of *ARHGAP35*, *RASA1*, *TJP1* and *TJP2* in normal lung tissue vs LUAD. (A)** *ARHGAP35*, *TJP1* and *TJP2* but not *RASA1* exhibit reductions in transcript levels in 524 LUAD samples relative to 59 samples of corresponding normal lung tissue. p = 5.889e-12, 0.2092, 2.185e-23, 5.392e-32, respectively. (**B**) Positive correlation between TJP2 and TJP1 expression levels in LUAD samples. Values are presented as VST transformed data by DESeq2.
